# ZNFX1, an immunoregulatory RNA helicase and E3 ubiquitin ligase, assembles into pleiomorphic polymers

**DOI:** 10.1101/2025.10.14.682297

**Authors:** Katerina Naydenova, Thomas Mund, Matthew C. J. Yip, Ceara M. Harper, Kendra E. Leigh, Jacqueline Hankinson, Keith B. Boyle, Alexander Heatley, Elsje G. Otten, Valeria Lulla, Yorgo Modis, Felix Randow

## Abstract

ZNFX1 is an SF1-family RNA helicase essential for innate immunity. Patients with ZNFX1 mutations experience recurrent infections, yet the underlying mechanism remains unclear. We determined cryo-EM structures of ZNFX1 in apo and RNA-bound forms, revealing auto-inhibition of the helicase through a regulatory insertion occluding the RNA-binding groove. ZNFX1 also functions as a bi-catalytic E3 ubiquitin ligase, containing an RZ-finger homologous to RNF213 and a previously unidentified Miz-like domain that catalyze ubiquitylation independently or cooperatively, with activity enhanced by ubiquitin chains. Patient mutations demonstrate that the helicase, E3 ligase, and rigid zinc-finger stalk connecting them are required for function. ZNFX1 can assemble into structured, pleiomorphic polymers via multivalent protein-protein interactions, revealing a mechanism that may facilitate RNA sequestration in stress granules and antiviral activity. These findings establish ZNFX1 as a multifunctional enzyme in innate immunity that couples RNA sensing to ubiquitin signaling and assembles into higher-order structures for signal amplification.

## Introduction

Innate immunity relies on a coordinated network of host factors to detect and respond to invading pathogens. Key components include pattern recognition receptors, signaling molecules, and effector proteins that regulate antiviral and antibacterial responses. Genetic defects, known as “inborn errors of immunity”, lead to severe immune deficiencies and render individuals susceptible to life-threatening and recurrent infections^1^. Studies of these genetic predispositions have provided critical insights into fundamental mechanisms of immune regulation and pathogen defense. They have also uncovered unexpected immune functions for previously unlinked genes^2^.

Mutations in the ZNFX1 (NFX1-type zinc finger-containing 1) gene were recently found to cause severe inborn errors of immunity^3–5^. Patients with deleterious homozygous or compound heterozygous ZNFX1 variants suffer from heightened predisposition to viral and bacterial infections and immune dysregulation, including intermittent monocytosis and hemophagocytic lymphohistiocytosis, frequently with fatal outcomes^3–5^. Additionally, ZNFX1 is among the most upregulated genes in SARS-CoV-2 infected cells^6^ and COVID-19 patients^7,8^.

While these studies revealed a direct involvement of ZNFX1 in primary immunodeficiency and anti-viral immunity, its specific role and mechanism of action remain unclear. ZNFX1 is an interferon-stimulated SF1-family RNA helicase, implicated in RNA sensing during viral infection^9^, and in host RNA stabilization during *Mycobacterium tuberculosis* infection^10^. ZNFX1 also contains an RZ domain, homologous to RNF213^11^, suggesting ZNFX1 may also function as an E3 ubiquitin ligase. A recent study, published while this manuscript was in preparation, confirmed that ZNFX1 is indeed an E3 ubiquitin ligase that, intriguingly, can ubiquitylate RNA *in vitro*^12^. To gain deeper insight into the molecular functions of ZNFX1, we determined its high-resolution structure and characterized its enzymatic activities *in vitro* and in cells.

Here we report the structure of human ZNFX1 in both ATP-bound and RNA-bound states, revealing a previously unrecognized auto-inhibitory mechanism where a negatively charged regulatory insertion plugs the RNA-binding groove of the helicase. We further demonstrate that the C-terminus of ZNFX1 contains a bi-catalytic E3 ubiquitin ligase, comprised of two non-canonical domains, an RZ finger and a Miz domain, that can catalyze ubiquitylation independently or cooperatively. Finally, we find that ZNFX1 assembles into pleiomorphic polymers via weak multivalent contacts, suggesting a role in RNA sequestration by higher-order ZNFX1 assemblies in cells. Together, these findings establish a structural and mechanistic framework for how ZNFX1 integrates RNA helicase and ubiquitin ligase activities for a novel type of antiviral defense strategy.

## Results

### Structure of human ZNFX1

To obtain insight into the molecular mechanisms governing ZNFX1 activity, we purified human ZNFX1 protein and determined its structure by cryogenic electron microscopy (cryo-EM) (**Figure 1A–C**, **S1A**, **Table S1**). Based on sequence homology, ZNFX1 had been predicted to be an SF1-family helicase. Indeed, its structure contains a typical SF1 helicase fold, with two characteristic RecA-like lobes (**Figure S2A**), expected to couple ATP hydrolysis to nucleic acid translocation. The N-terminal helicase lobe contains two regulatory insertions: a beta-barrel, commonly termed 1B, and a longer insertion, 1C, which is only partially resolved in the cryo-EM map (**Figure 1B–C, S2A**). Based on the cryo-EM structure, we annotated the domain features for the helicase domain of ZNFX1 (**Figure 1A)**. An additional small N-terminal domain with an armadillo-type fold comprising eight alpha helices is also resolved in the cryo-EM map, despite being flexibly tethered to the helicase (**Figure 1A–C**). Surprisingly, the NFX1-type zinc fingers, previously annotated by sequence homology, extend at a fixed angle from the helicase as a rigid linker, rather than a flexible ‘pearls-on-a-string’ arrangement, and are well resolved in the cryo-EM map to approximately residue 1430 (**Figure 1A–C**). The remaining, circa 500-residue C-terminal fragment is not resolved in the cryo-EM map, indicating high internal flexibility as it could not be resolved even by local refinement strategies.

**Figure 1.**
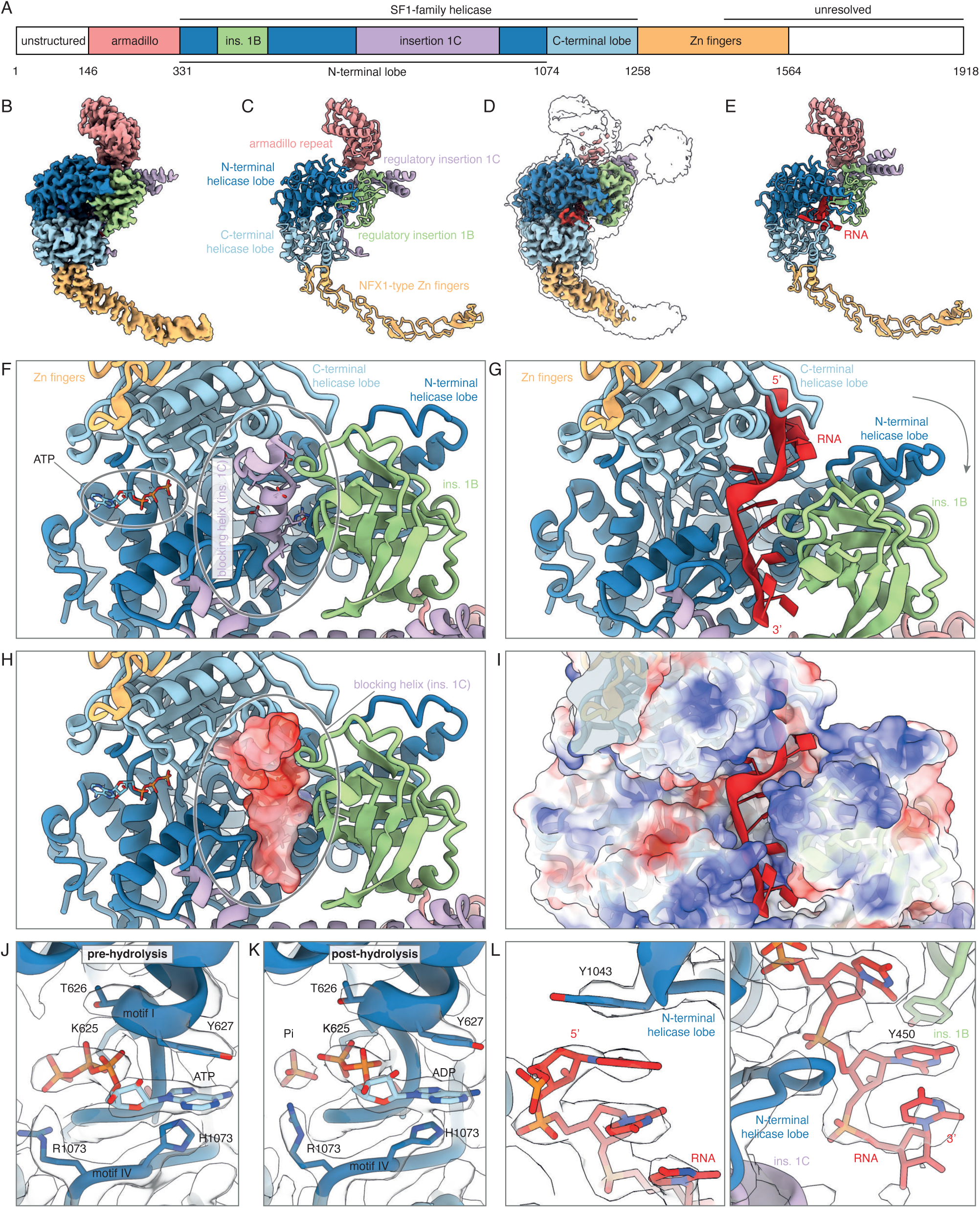
Structural overview of ZNFX1. (A) Domain annotation of human ZNFX1. (B) Cryo-EM structure and model (C) of human ZNFX1. (D) Cryo-EM structure and model (E) of human ZNFX1, bound to RNA. Domains are color-coded in all panels as follows: armadillo repeat, pink; N-terminal helicase lobe, dark blue; C-terminal helicase lobe, light blue; regulatory insertion 1B, green; regulatory insertion 1C, violet; NFX1-type Zn fingers, orange. (F) Zoomed in view of the blocking helix from regulatory insertion 1C occluding the RNA-binding groove of the ZNFX1 helicase domain, and (G) the same view of the groove when RNA is bound. (H) The same view as in (F), with the surface of the “plug” helix colored by electrostatic charge. (I) The same view as in (G), with the surface of ZNFX1 colored by electrostatic charge. (J, K) Zoomed in view of the cryo-EM density map and model of the (J) pre-and (K) post-ATP hydrolysis states of the ZNFX1 in the helicase domain. (L, M) Cryo-EM density map and model of RNA bound to the helicase domain of ZNFX1, zoomed in views at around the 5’ end (L), and at the 3’ end (M) of the resolved RNA strand in the structure.

### Structure and regulation of the ZNFX1 helicase domain

To investigate the molecular mechanism of the ZNFX1 helicase, we determined its structure in the presence of ATP and RNA, added separately and together (**Figure 1B–E, S1–S2**). In the presence of ATP alone, the ZNFX1 helicase domain adopts a semi-closed conformation containing unhydrolyzed ATP (**Figure 1J**). The nucleotide is bound in the cleft between the conserved Walker A ATP-binding motif I and the Walker B ATP-hydrolysis motif II (**Figure S2A**). The Walker B motif in ZNFX1 (residues 1006 – 1009) features a deviant EE_1007_AA sequence, distinct from the canonical DExx motif in SF1 helicases^13^, with E1007 identified as the catalytic residue required for ATP hydrolysis.

As revealed by the cryo-EM structure (**Figure 1J**), ZNFX1 does not hydrolyze ATP at an appreciable rate in the absence of RNA, a hallmark of auto-inhibited helicases^14^. To understand the mechanism of ZNFX1 auto-inhibition, we compared structures determined in the absence and presence of RNA. In the RNA-bound, ATP-free structure (**Figure 1D–E**), a single-stranded nucleic acid occupies an RNA-binding groove between the two RecA-like helicase lobes. In the absence of nucleic acid, the same groove is occluded by a short alpha helix (residues 828 – 836, part of the 1C regulatory insertion), suspended on flexible loops at both ends (**Figure 1F–G**). Notably, this “plug” helix is a conservation hotspot within the otherwise variable insertion (**Figure S3**) and is exceptionally negatively charged (**Figure 1H**), mimicking the RNA backbone.

RNA binding is sufficient to displace the “plug” helix, which becomes unresolved in the cryo-EM map (**Figure 1G**), and this also shifts the regulatory insertion 1B to clamp in the RNA (**Figure 1F–G**). We thus propose ZNFX1 is auto-inhibited through binding of the “plug” helix to the RNA binding groove between the two helicase lobes, representing a previously unrecognized mode of auto-inhibition in SF1 helicases. RNA contacts the positively charged groove mainly via its phosphate backbone (**Figure 1I**), except for two notable stacking contacts: between Y1043 and the most 5’ terminal resolved RNA base, and between Y450 in insertion 1B and an RNA base near the 3’ exit site (**Figure 1L**). These stacking contacts likely mediate the clamping mechanism that couples ATP hydrolysis to RNA translocation.

Finally, we determined the structure of ZNFX1 in the presence of both ATP and RNA. In the resulting structure, no RNA density was detected. The RNA-binding groove, instead of containing RNA, is occluded by the “plug” helix, as in the structure determined without added RNA. The nucleotide binding pocket, however, is occupied by ADP and Pi, the products of ATP hydrolysis (**Figure 1K**), demonstrating that ATP hydrolysis was stimulated by RNA, followed by its dissociation. We propose this structure represents the post-hydrolysis state of the helicase.

The closest structural match to the helicase domain of ZNFX1, identified using PDBeFold^15^, is UPF1, a 3’-directed SF1-family helicase involved in nonsense-mediated RNA decay^16^ (**Figure S2**). The main distinctive feature of the ZNFX1 helicase domain is the unusually large 1C insertion (residues 688 – 978), far exceeding that of UPF1. This region shows no structural homology to any previously determined structure. The closest sequence homolog for the insertion is found in the fission yeast (*Schizosaccharomyces pombe*) helicase HRR1, involved in RNAi^17^, which remains uncharacterized functionally and may also have a similar mechanism for auto-inhibition. Interestingly, ZNFX1 is also reported to participate in RNAi in *C. elegans*^18^.

In summary, our cryo-EM structures demonstrate that ZNFX1 is an auto-inhibited 5’-3’ RNA helicase of the SF1 family. It exhibits a distinctive regulatory insertion: a negatively charged helix within insertion 1C that occupies the RNA-binding groove, providing a novel mechanism for auto-inhibition that may also apply to other helicases, such as the RNAi-mediator HRR1.

### Patient mutations inactivate the helicase domain of ZNFX1

We used the cryo-EM structures to investigate the impact of three deleterious mutations in the ZNFX1 helicase domain (V469L, L1051P and R1223Q^4,5^), previously identified in patients with recurrent infections (**Figure 2A–D**, **Table S2**). All three residues are >98% conserved across chordates (**Figure S3**). Although seemingly conservative, the leucine-to-valine mutation (V469L) in regulatory insertion 1B likely destabilizes the entire insertion, as V469 is tightly packed against surrounding hydrophobic residues (**Figure 2B**), leaving insufficient space to accommodate the bulkier leucine side. The side chain of L1051, located in the N-terminal helicase lobe, is similarly oriented towards the hydrophobic protein core (**Figure 2C**), and its mutation to proline, a smaller, cyclic residue, likely disrupts local folding. Finally, the conserved R1223 is part of the consensus sequence motif VI in SF1 helicases^19^ (**Figures S2–S3**), coordinating the gamma phosphate of bound ATP (**Figure 2D**). We conclude that all three patient mutations disrupt the helicase activity of ZNFX1 by destabilizing the regulatory insertion or helicase fold, or by impairing ATP binding and hydrolysis.

**Figure 2.**
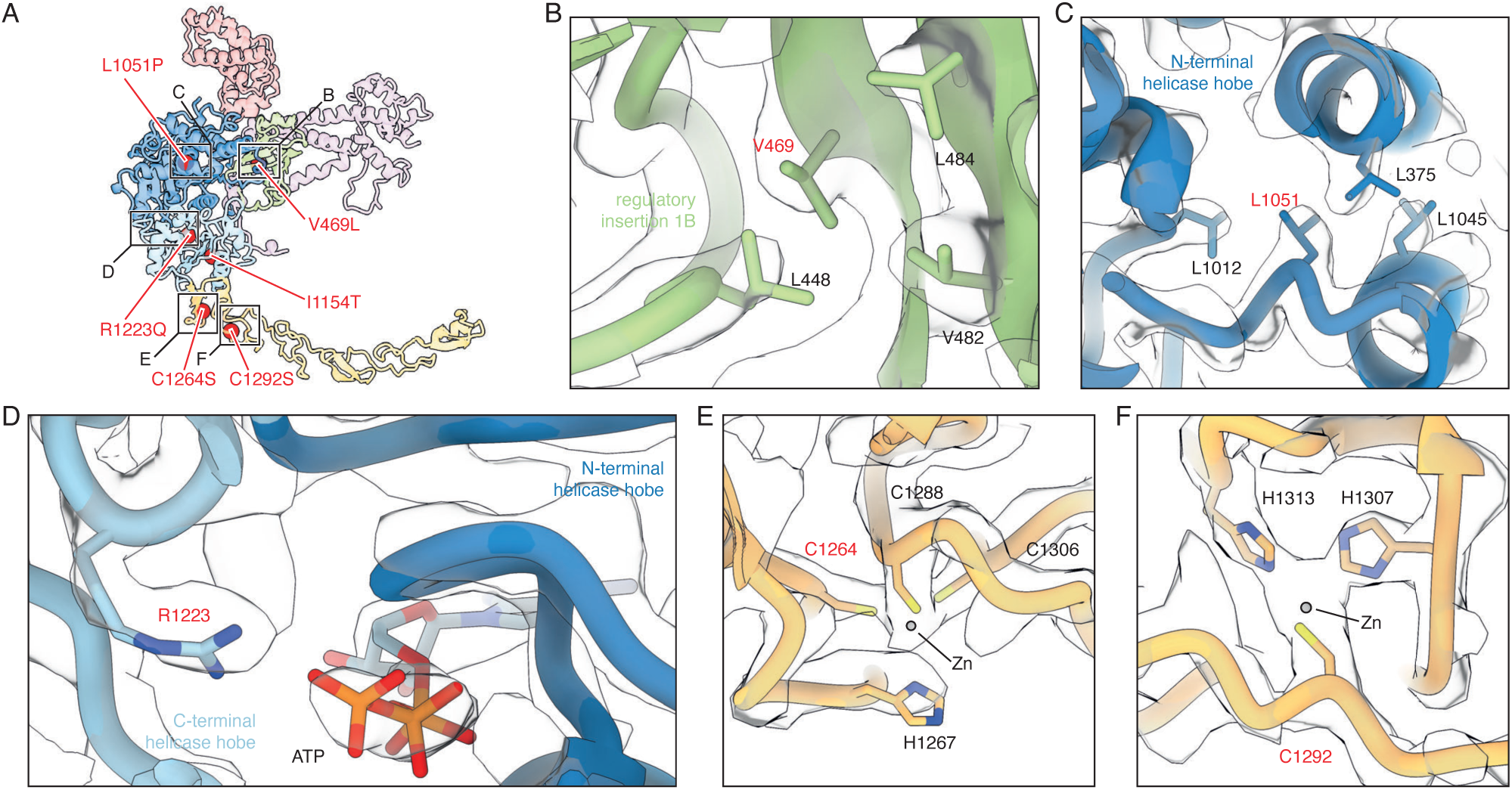
Patient mutations mapped to the structure of ZNFX1. (A) Overview of the structure of ZNFX1, with the sites of deleterious patient mutations (**Supplementary Table S2**) highlighted as red spheres. The black boxes outline the approximate regions, which are zoomed into in the following panels. (B-F) Zoomed in view of the ZNFX1 structure around the deleterious patient mutations, with the cryo-EM density map overlayed as transparent surface.

### The ZNFX1 NFX1-type Zn fingers form a rigid stalk

Next, we mapped two deleterious mutations (C1264S and C1292S), previously identified in patients with recurring infections, to the Zn finger domain of ZNFX1 (**Figure 2E–F**, **Table S2**). Both residues are strictly conserved across species (**Figure S3**) and are well resolved in the cryo-EM maps, where they coordinate Zn^2+^ ions. Substitutions with serine likely disrupt Zn^2+^ binding in the first and third Zn finger, respectively, causing domain unfolding and destabilizing the rigid Zn finger stalk that links the helicase to the C-terminus. While the exact function of the Zn fingers stalk remains unknown, the patient mutations highlight its functional importance *in vivo*.

### ZNFX1 forms pleiomorphic helical assemblies in vitro

A potential function of the zinc fingers was revealed during early cryo-EM analysis, when we observed helical filaments in the electron micrographs of human ZNFX1 protein (**Figure S1A**), purified from insect cells, and used at a relatively high concentration (2–4 mg/mL). Two cryo-EM specimens, prepared from independently purified batches (***Methods***), revealed distinct filament morphologies. One specimen contained short, helical filaments that remained dispersed, whereas the other contained bundles of long, nearly straight fibrils forming twisted sheets (**Figure 3A–F**).

**Figure 3.**
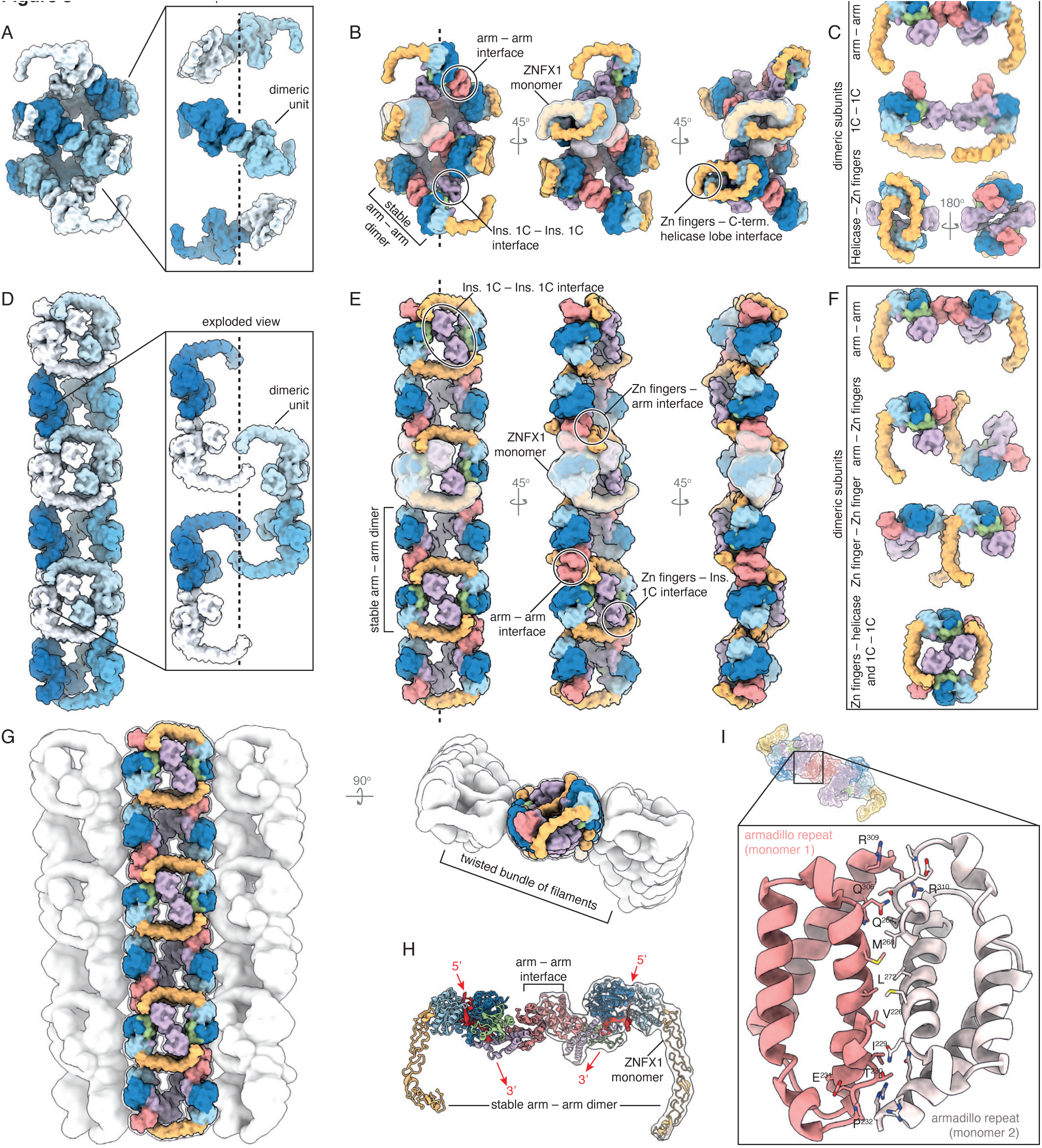
ZNFX1 forms pleiomorphic polymers. (A–F) Cryo-EM models of two types of ZNFX1 polymers: a short helical filament (A–C), and a bundle of long twisted filaments (D–F). The filaments are colored by ZNFX1 monomer in (A, D), or by domain as in Figure 1 in (B–C, E–F). The insets show an exploded view of a piece of each filament, made up of repeating ZNFX1 dimer subunits. (C, F) Dimeric ZNFX1 units that compose each type of filament, showcasing the relevant inter-protein contacts. (G) The low-resolution cryo-EM map of the bundle of twisted filaments (white) is overlayed on the model of a single filament. In (B, E) one monomer is outlined for clarity, and the inter-protein interactions forming the filaments are highlighted. The highlighted common armadillo-domain mediated dimer, also resolved in solution is shown in (H), with the RNA strand directions indicated by arrows. (I) Zoomed in view of the armadillo domain-mediated C2-symmetric dimerization interface.

The repeating unit in both polymeric forms is a C2-symmetric ZNFX1 dimer, mediated by homophilic armadillo domain contacts (**Figure 3A, D**). In the short filaments consecutive dimers interact via the Zn-finger stalk of one monomer wrapping around the C-terminal lobe of the helicase domain of another monomer in the next dimer. Additional homophilic contacts through regulatory insertion 1C are formed between non-consecutive dimers (i and i+2) (**Figure 3B–C**). The long fibrils are stabilized by a different set of inter-dimer interactions: homophilic contacts between pairs of zinc fingers, between pairs of insertions 1C, and heterophilic contacts of zinc fingers with armadillo and helicase domains (**Figure 3E–F**). Sequential dimers are related by a ∼180° rotation around the helical symmetry axis and a translation of approximately one monomer along the same axis. Unlike the short filaments, the long fibrils formed parallel, twisted bundles or sheets (**Figure 3G**). Inter-fibril contacts, established between helicase domains, are weaker than fibril-forming contacts, resulting in more disordered assemblies.

In both polymeric forms, all ZNFX1 molecules orient with the 3’ exit of the helicase towards the filament interior, making it unlikely that a single RNA molecule traverses the assembly axially. The ability of ZNFX1 to form distinct polymeric arrangements based on the weak, multi-valent protein-protein contacts revealed here may underlie the nucleic acid-dependent activation and clustering of ZNFX1 reported by others^12,20^. We propose that different ZNFX1 assemblies may form in cells to sequester viral RNAs, organize host RNAs into stress granules, or store inactive ZNFX1 protein. We hypothesize that ZNFX1 can dynamically switch between the different polymeric forms in response to cellular cues.

### The ZNFX1 armadillo domain is a homodimerization interface

A defining feature of all ZNFX1 polymers is the C2-symmetric dimer formed by two opposing armadillo-repeat domains (**Figure 3H**). The same dimer was also observed as a minor species in other cryo-EM specimens, where most ZNFX1 remained monomeric. We therefore hypothesize that armadillo-armadillo contacts (**Figure 3I**) represent the principal interaction driving polymer formation. The high-resolution structure of this interface (**Figure 3I**) will guide mutational analysis to test the functional significance of ZNFX1 dimerization *in vivo*.

### The RZ domain contains a conserved ubiquitin-binding motif required for catalytic activity

The C-terminal fragment of ZNFX1 (residues 1434–1918) was not resolved in any cryo-EM map (**Figure 1**), suggesting substantial flexibility relative to the helicase domain and the rigidly attached NFX1-type zinc fingers. We therefore investigated this region further through a combination of structure prediction and biochemical assays.

We previously reported that the C-terminal fragment of ZNFX1 encodes a novel E3 ubiquitin ligase domain, homologous to RNF213, thus defining the RZ (RNF213–ZNFX1) finger^11^. The RZ domain in ZNFX1 can be traced back to unicellular eukaryotes, e.g. choanoflagellates^11^. Our cryo-EM structure of RNF213 revealed that the RZ domain adopts a β-sandwich fold, preceded by a short and flexibly hinged α-helix. The fold is stabilized through a Zn²⁺ ion coordinated by conserved Cys and His residues^21^, and contains flexible, unresolved loops predicted to position a pair of catalytic residues, C4516 and H4537, in exact proximity. AlphaFold structure predictions indicate that the ZNFX1 RZ domain shares this architecture (**Figure 4A**), with C1849, H1853, C1869, and C1872 forming the metal-binding site, and C1860 and H1881 corresponding to the nucleophile and general base, respectively, as previously identified in RNF213^21^.

**Figure 4.**
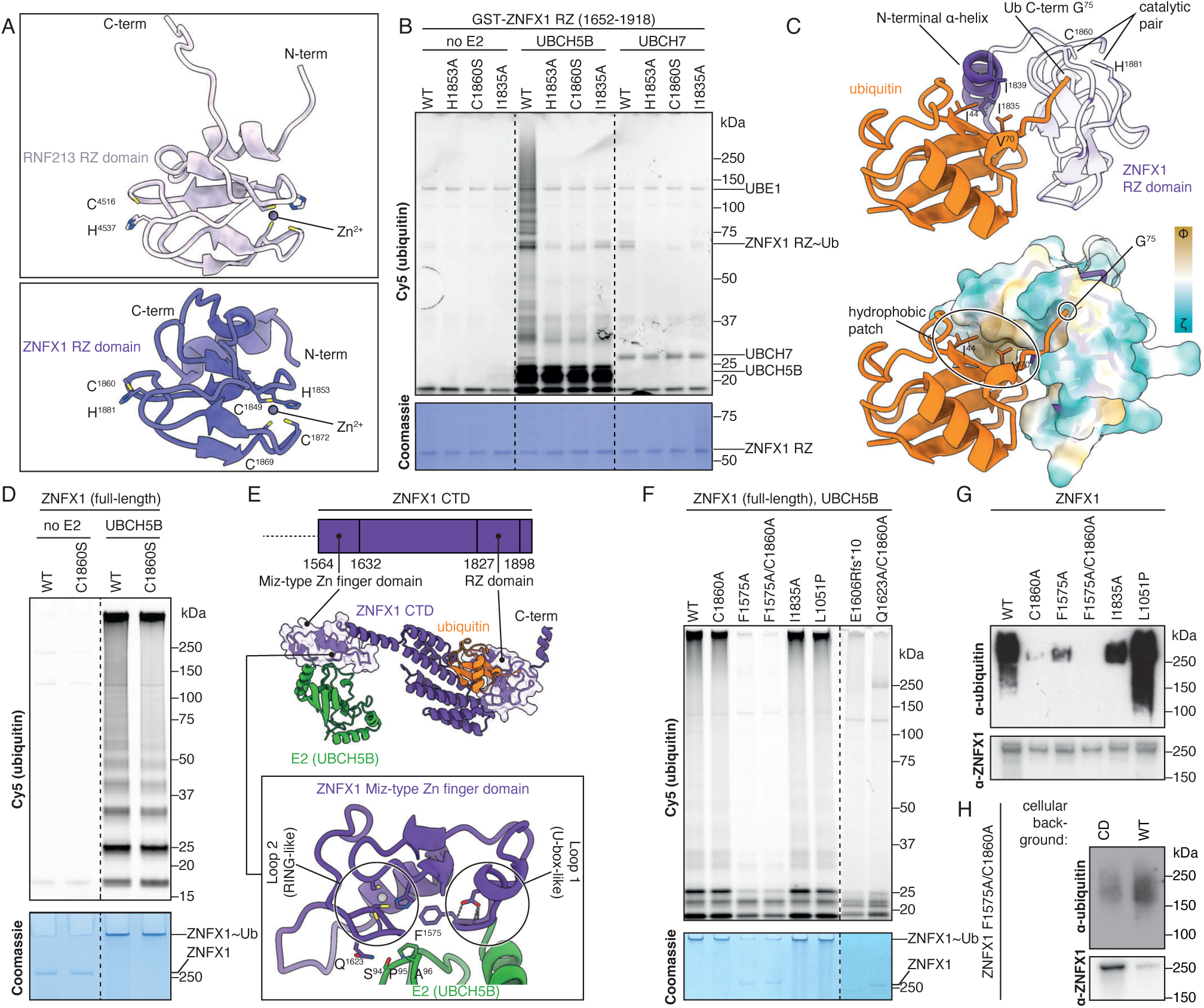
ZNFX1 is a dual E3 ubiquitin ligase utilizing a Miz and RZ domains for full activity. (A) Ribbon diagrams of predicted RZ domain structures of RNF213 (top) and ZNFX1 (bottom) indicating the coordinated Zn^2+^ ions and the respective predicted catalytic cysteine/histidine pairs. (B) E3 ubiquitin ligase catalytic activity of the ZNFX1 RZ domain fragment (see also domain diagram in (E)) expressed in bacteria (WT and various point mutations as indicated) with UBCH5B, UBCH7 or no E2 enzyme control, monitored *in vitro* by Cy5-ubiquitin fluorescence and Coomassie staining of SDS-PAGE. (C) Structure prediction of the ZNFX1 RZ domain (purple/white) in complex with ubiquitin (orange), highlighting the C-terminal glycine of ubiquitin, the catalytic cysteine/histidine pair (C1860 and H1881), and the predicted binding interface of the major hydrophobic patch of ubiquitin with the N-terminal alpha helix of the RZ domain. Lower panel, RZ domain surface colored by hydropathy: Φ, hydrophobic (yellow); ζ, hydrophilic (blue). (D) *In vitro* ubiquitylation assays with full-length human ZNFX1 expressed in insect cells (WT or catalytic C1860S mutant) with or without UBCH5B, monitored by Cy5-ubiquitin fluorescence and Coomassie staining of SDS-PAGE. (E) Structure prediction of the C-terminal fragment of ZNFX1 (purple), bound to an E2 enzyme (UBCH5B, green) and to ubiquitin (orange). Inset, predicted structure of the newly identified Miz-type domain of ZNFX1 in complex with UBCH5B, highlighting the U-box/RING-like domain and the E2 interaction interface. (F) *In vitro* ubiquitylation assays as in (D) using various ZNFX1 mutants, expressed in human cells, as indicated. (G) Anti-ubiquitin immunoblot of ZNFX1 proteins purified from human cells demonstrating their ubiquitylation status prior to being used in (F). Anti-ZNFX1, loading control. (H) Anti-ubiquitin immunoblot of ZNFX1 F1575A/C1860A protein, purified from human cells co-expressing wild-type (WT) or E3 catalytic dead mutant (CD, F1575A/C1860A) ZNFX1 with a different tag. Anti-ZNFX1, loading control.

To test whether the RZ domain fragment of human ZNFX1 functions as an E3 ubiquitin ligase, we performed *in vitro* ubiquitylation reactions. The ZNFX1 RZ fragment exhibited auto-ubiquitylating activity (**Figure 4B, S4C**), with both Zn^2+^ coordination (H1853) and the putative catalytic cysteine (C1860) required for this activity. Unlike RNF213^22^, the ZNFX1 RZ fragment preferentially utilizes the E2 enzyme UBCH5B over UBCH7 (**Figure 4B**).

To obtain further mechanistic insight into the reaction mechanism of the RZ domain, we investigated possible interactions with ubiquitin and E2 ubiquitin-conjugating enzymes. AlphaFold structure predictions suggest that the N-terminal alpha helix of the RZ domains in both RNF213 and ZNFX1 (**Figure 4C**, **Supplementary Figure S4A**) binds the major hydrophobic patch of ubiquitin (I44, V70). This binding mode positions the ubiquitin C-terminal glycine near the catalytic cysteine (ZNFX1 C1860 or RNF213 C4516), suggesting the interaction is required for RZ domain activity. Indeed, mutating a conserved residue (I1835A) on the predicted ubiquitin-binding helix abolished catalytic activity (**Figure 4B**). We conclude that ubiquitin binding by the RZ domain is required for its E3 ligase activity.

### ZNFX1 is a bi-catalytic E3 ubiquitin ligase comprised of two non-canonical E3 ligase domains

Surprisingly, mutations in the RZ finger (C1860S, C1860A, or I1835A) that abrogate activity of the isolated domain, did not abolish E3 ubiquitin ligase activity in full-length ZNFX1 *in vitro* (**Figure 4D**), indicating the presence of a second, hitherto unknown E3 ligase domain in ZNFX1. To identify this alternative domain, we compared the predicted structure of the unresolved ZNFX1 C-terminal fragment, excluding the NFX1-type Zn fingers and RZ domain, to previously determined structures using PDBeFold^15^. The search revealed close resemblance between ZNFX1 residues 1566 – 1632 and the U-box domain of the yeast E4 ubiquitin ligase Ufd2^23^. U-box domains bind E2 ubiquitin-conjugating enzymes and function as E3 and E4 ubiquitin ligases. Their cross-brace topology resembles that of RING domains, but lacks the metal-coordinating core^24^. However, unlike either RINGs or U-boxes, the homologous domain in ZNFX1 can accommodate a single zinc ion (in site II), with site I stabilized by a hydrogen bond network instead, as in U-box domains **(Figure 4E, S4B**). The loop-forming residues are phylogenetically conserved **(Figure S3**). Such a “hybrid” U-box/RING-like domain has otherwise been found exclusively in SUMO E3 ligases of the Siz/PIAS family, where it is known as SP-RING or Miz-type RING; we refer to it as Miz domain hereafter. We demonstrate that the Miz domain of ZNFX1 is an active ubiquitin ligase (**Figure 4F, S4D**), even in the absence of catalytic activity in the RZ domain, thus constituting the first example of a Miz domain with E3 ligase activity for ubiquitin. Interestingly, *C. elegans* ZNFX1 lacks the RZ domain entirely but likely retains ubiquitin ligase activity through its Miz domain. The patient-derived truncated variant ZNFX1 E1606Rfs*, lacking both RZ and Miz domains, is completely inactive in ubiquitylation assays **(Figure 4F**), whereas the patient-derived helicase mutant L1051P retains auto-ubiquitylating activity comparable to that of wild-type ZNFX1 **(Figure 4F**). Taken together, our results demonstrate that ZNFX1 contains two distinct non-canonical E3 ligase domains, an RZ finger and a Miz domain, and that loss of E3 ligase activity predisposes to infection.

### Interaction of the ZNFX1 Miz domain with ubiquitin-conjugating E2 enzymes

The human ZNFX1 Miz domain lacks the typical arginine ‘linchpin’ residue^25^ that facilitates ubiquitin transfer from the E2 enzyme to the substrate. Instead, a glutamine (Q1623 in human ZNFX1) is found at this position in most species. A sub-optimal glutamine ‘linchpin’ is also found in other active RING domains, for example TRIM72^26^. Mutation of this residue to alanine, Q1623A, in the context of an inactive RZ domain (C1860A), completely abolished E3 ubiquitin ligase activity *in vitro* (**Figure 4F**). We therefore conclude that the ZNFX1 Miz domain facilitates ubiquitin transfer by a ‘linchpin’-based mechanism.

Based on structure prediction and comparison with previously determined RING– and U-box– E2 complexes, we disrupted the predicted E2 binding site in the Miz domain by introducing the F1575A mutation. This mutation, combined with RZ-domain inactivation (C1860A), caused complete loss of ZNFX1 E3 activity (**Figure 4F, S4B, E**). To further investigate the ability of ZNFX1 to function with specific E2 enzymes, we screened a library of human E2 enzymes in an *in vitro* ubiquitylation reaction. ZNFX1 exhibited strong preference for the UBCH5 family (Ube2D1–4) and detectable activity with Ube2E1–3. This preference is explained by the complementary SPA motif in loop 7 of preferred E2 enzymes, whereas the bulkier KPA motif in the equivalent position of UBCH7 cannot be accommodated by the ZNFX1 Miz domain. The determinants of RZ-domain preference for UBCH5B over UBCH7, however, remain unclear. We also noticed that in the SUMO E2 UBE2I (Ubc9), the corresponding RPA motif is similarly incompatible with the ZNFX1 Miz domain (**Figure S5B–C**). Our results thus identify the ZNFX1 Miz domain as the first Miz-family member to function in the ubiquitin conjugation pathway.

### Bi-catalytic E3 ubiquitin ligase activity of ZNFX1 in vivo

We next compared the relative contributions of the Miz and RZ domains to ZNFX1 E3 ligase activity *in vitro* and in cells. We tested ZNFX1 C1860S, lacking the catalytic residue of the RZ domain, and ZNFX1 F1575A, defective in E2 binding via its Miz domain. With purified proteins *in vitro*, most auto-ubiquitylating activity originated from the Miz domain (**Figure 4F**) while ZNFX1 C1860S retained wild-type activity. In contrast, in *ZNFX1 KO* cells complemented with the relevant alleles, defects in either domain severely impaired auto-ubiquitylation (**Figure 4G**), revealing that both domains are required for E3 activity in cells. Ubiquitylation of ZNFX1 was abolished when both domains are mutated (**Figure 4G**), demonstrating that ZNFX1 ubiquitylation originates from intrinsic E3 ligase activity. We therefore attribute the very low auto-ubiquitylating activity of E3 domain mutants (C1860A and F1575A) in cells to residual activity of their corresponding wild-type Miz and RZ domain. We conclude that the activity of wild-type ZNFX1 is much greater than the sum of the activities of its individual E3 domains (**Figure 4G**). We therefore propose that ZNFX1 is a bi-catalytic E3 ubiquitin ligase, where maximal efficiency is achieved when the two domains cooperate to transfer ubiquitin in a linear cascade, likely from the Miz domain-bound E2 to the RZ and then to the final substrate, in agreement with another recent study^12^.

In support of the bi-catalytic mechanism, patient-derived truncations (E1727fs*11 and V1788fs*6) that lack an RZ finger but retain the Miz domain confer susceptibility to infection **(Figure 4E)**. In contrast, the helicase-domain mutation L1051P does not impair auto-ubiquitylating activity in cells (**Figure 4G**).

Considering the ability of ZNFX1 to polymerize, its auto-ubiquitylating activity could manifest *in cis* or *in trans.* To test for *trans*-autoubiquitylation, we co-expressed catalytically inactive ZNFX1 F1575A/C1860A and wild-type ZNFX1, serving as substrate and enzyme, respectively. Co-expression of wild-type ZNFX1 markedly enhanced the ubiquitylation of inactive ZNFX1 and reduced its protein level, thus revealing the occurrence of trans-autoubiquitylation and ubiquitylation-dependent protein turnover (**Figure 4H**).

### The E3 ubiquitin ligase activity of ZNFX1 is allosterically enhanced by extended ubiquitin ***chains***

To further investigate how ZNFX1 E3 ligase activity is modulated, we tested its response to ubiquitin chains. ZNFX1 catalytic activity was strongly enhanced *in vitro* by linear (M1-linked) or K63-linked tetra-ubiquitin chains, weakly by linear di-ubiquitin, but not by K48-linked tetra-ubiquitin (**Figure S5E–G**). This suggests that activation requires a minimal ubiquitin chain length and extended chain topologies, as found in M1-and K63-linked but not K48-linked chains.^27^ Ubiquitin chains stimulate full-length ZNFX1 (**Figure S5E**) and C-terminal fragments lacking armadillo and helicase domains (**Figure S5F**), including mutants with inactive RZ domains (**Figure S5G**). We conclude that ubiquitin-chain mediated potentiation of ZNFX1 E3 activity occurs via the Miz domain, independent of helicase activity, RZ activity, armadillo-mediated dimerization, or polymerization. The activation mechanism is likely allosteric, as the supplied ubiquitin chains were not consumed in the assay (**Figure S5E**), indicating they act as regulatory factors not substrates.

## Discussion

Mutations in ZNFX1 predispose individuals to recurrent viral and bacterial infections. Despite its clinical importance, the molecular basis of ZNFX1 function has remained unclear. Through cryo-EM, we defined the distinct architecture of ZNFX1, which integrates RNA helicase and E3 ubiquitin ligase activities with an intrinsic propensity to assemble into pleiomorphic polymers. These structural features support a coordinated mechanism that couples RNA recognition and activation of E3 ligase activity. Our findings provide a mechanistic framework to understand how patient-derived mutations disrupt ZNFX1 function and compromise host defense.

ZNFX1 encodes a canonical SF1 RNA helicase domain that is autoinhibited by a previously unrecognized, negatively charged ‘plug’ helix. In addition, the protein contains a distinctive ’bi-catalytic’ E3 ubiquitin ligase module comprised of two non-canonical catalytic elements: (i) a Miz-like U-box/RING ‘hybrid’, and (ii) an RZ finger, a domain found only in ZNFX1 and in RNF213 among human proteins. ZNFX1 assembles into pleiomorphic polymers, nucleated by armadillo domain-mediated dimerization and further stabilized through weaker, multivalent contacts. Mapping deleterious patient mutations onto these structures reveals that helicase activity, E3 ligase activity, and the rigid Zn finger stalk that promotes polymerization are all essential for the physiological function of ZNFX1 in protection against recurrent infection.

The bi-catalytic E3 module in ZNFX1 resembles RBR E3 ligases, with their two-step catalytic mechanism, in which ubiquitin is first transferred from a RING1-bound E2 enzyme to a catalytic cysteine in RING2, before reaching the substrate. Such sequential transfer from Miz-bound E2 to C1860 in the RZ finger appears to be the primary mechanism for ZNFX1 ubiquitylation in cells, consistent with mutations in either the Miz or the RZ domain largely eliminating activity. *In vitro*, in contrast, direct ubiquitin transfer from the Miz domain to the substrate predominates, revealing a reaction mechanism similar to that of canonical RING-domain E3 ligases. ZNFX1 can therefore switch between a RING-like, one-step mode and an RBR-like, bi-catalytic mechanism. It is tempting to speculate that in cells, assembly of ZNFX1 into polymeric structures positions the RZ domains as the preferred reaction partners of Miz-bound E2 enzymes, thereby favoring RBR-like catalysis. This mechanism may require cooperation between Miz and RZ domains from different protomers, consistent with evidence for trans-autoubiquitylation in cells. We propose that cellular context and stimuli, by controlling ZNFX1 filament formation and facilitating *trans* interaction between Miz and RZ domains, can tune the balance between these two catalytic modes to affect the ubiquitin signaling outcome.

An additional layer of regulation was recently revealed by two studies (one published, the other on a preprint server)^12,20^ which showed that RNA binding enhances ZNFX1 E3 ubiquitin ligase activity. This activation drives formation of ubiquitin-coated nucleoprotein particles that cluster single-stranded RNA and restrict viral replication^12,20^. Our cryo-EM structures provide mechanistic insights into how RNA activates ZNFX1 and how ZNFX1 forms nucleoprotein particles. First, we reveal that the structural basis for the previously unexplained RNA-mediated allosteric activation of the ZNFX1 E3 ubiquitin ligase activity lies in the displacement of the ‘plug’ helix by incoming RNA. Displacement of this helix represents the principal structural rearrangement upon RNA binding; it permits helicase function by granting RNA access to the helicase core. The displaced helix itself could also participate in activating the E3 ligase, suggesting an elegant unifying mechanism that coordinates helicase and ligase functions. Further experiments are required to test this hypothesis. Second, by comparing the structures of two distinct ZNFX1 polymers, we demonstrate that ZNFX1 has the propensity for multiple, weak self-interactions – a hallmark of proteins that form dynamic, reversible assemblies, well suited for stress granule formation and nucleic acid entrapment. In our hands, ZNFX1 dimers and higher-order assemblies formed readily even in the absence of nucleic acids, suggesting that protein-protein interactions alone are sufficient for polymerization. Third, the filamentous ZNFX1 assemblies provide a simple geometric explanation for RNA entrapment. As all ZNFX1 monomers are oriented with their 5’ helicase exit sites facing the filament exterior, RNA traversing more than one ZNFX1 monomer must wrap around ZNFX1 and re-thread into the interior of the filament. If this RNA is ubiquitylated by ZNFX1, as suggested by one of the recent studies^12^, collective helicase activity in the oligomer would eventually bring the modified base in contact with another helicase domain, thereby stalling further translocation.

The recent discoveries of RNA ubiquitylation by ZNFX1 *in vitro* and of LPS ubiquitylation by RNF213 during bacterial infection pose the question of what endows these two RZ domains with their special ability to ubiquitylate non-proteinaceous substrates. We propose that the ubiquitin-binding site in RZ domains, identified here, contributes to enabling ester-linked ubiquitylation. Ubiquitin binding by the RZ domain appears analogous to the way specialized E2 enzymes, such as UBE2J2, bind the major hydrophobic patch of the covalently bound donor ubiquitin, thereby causing a rearrangement of the E2 active site that enhances the reactivity of the E2-Ub thioester to facilitate attack by weaker nucleophiles^28^. In addition, a nearby histidine activates a substrate hydroxyl group by general base catalysis, a feature also shared with RZ domains.

Formation of K63/M1-linked ubiquitin chains is markedly induced in cells stimulated by inflammatory cytokines. Similar to their roles in activating NFκB and JNK signalling^29^, we discovered that they also stimulate ZNFX1 E3 activity, potentially contributing to ZNFX1 activation during viral infection. The potentiation of Miz domain activity by extended ubiquitin chains suggests a feed-forward loop, as parallel work^20^ demonstrates that the Miz domain itself preferentially synthesizes K63-linked chains. Further elucidation of this activation mechanism, which resembles the allosteric activation of HOIL-1L by K63/M1-linked ubiquitin chains^30^, will be important. Based on our ZNFX1 fragment constructs, we identify candidate structural determinants for this allosteric mechanism: the flexible, C-terminal portion of the zinc fingers, the Miz domain, and the helical bundle linking Miz and RZ domains. Since certain zinc fingers bind ubiquitin chains, while others interact with nucleic acids, it is tempting to speculate they could even confer specificity for ubiquitylated nucleic acid. The ubiquitin chain-mediated feed-forward loop likely explains the strong Miz domain activity observed *in vitro* at high local concentrations of ubiquitin and ubiquitylated substrates. A similar scenario may occur in ZNFX1-rich cellular compartments, such as RNA-containing stress granules. Since the ubiquitin chain-mediated feed-forward loop proceeds independently of RZ domain activity, this regulatory mechanism may also operate in evolutionarily distant context, e.g. in *C. elegans*, where ZNFX1 lacks an RZ domain yet traps RNAs in nucleoprotein granules^18,31^.

Our work has revealed that ZNFX1 activity is controlled through at least four mechanisms: RNA-induced de-inhibition, ubiquitin chain-mediated E3 activation, switching to a bi-catalytic E3 mechanism, and polymerization. Protein clustering for signal amplification is a prevailing theme in inflammatory signaling, illustrated by MDA5 filament formation on viral nucleic acids^32^, MAVS oligomerization on mitochondria^33^, cGAS filament formation on exogenous DNA^34^, or RNF213 polymerization on the surface of pathogens^21^, among numerous other examples. We have now shown that ZNFX1 also conforms to this paradigm. The physiological and infection-related triggers and cellular compartments where ZNFX1 assemblies form – stress granules^3,35^, the mitochondrial surface^9^, or in the vicinity of a pathogen^35^ – remain to be further investigated. We predict that ZNFX1-rich structures in cells serve as ubiquitin signaling platforms, thereby linking the catalytic activities of ZNFX1 to broader inflammatory signaling.

## Methods

### Antibodies

The following primary antibodies were used in this work: anti-ubiquitin (Life Sensors, VU-1, 1:1,000), anti-ZNFX1 (Abcam, ab179452, 1:1,000). The secondary antibodies used were from Dako (HRP-conjugated goat anti-mouse and anti-rabbit).

### Cell culture

Cells were grown in IMDM supplemented with 10% FCS at 37°C in 5% CO_2_. All cell lines tested negative for *Mycoplasma* contamination.

### Generation of knockout cell lines

To make *ZNFX1*-knockout cells, oligonucleotide duplexes for the gRNA were phosphorylated with T4 PNK (New England Biolabs) and cloned into pSpCas9(BB)-2A-GFP (Addgene, PX458). HEK293ET cells were transfected, and single GFP-positive cells were sorted into 96-well plates 24 hours later. Resulting clones were screened for lack of ZNFX1 expression by immunoblotting, and disruption of the *ZNFX1* exon was verified by sequencing.

### Protein expression and purification

Short ZNFX1 fragments were expressed as N-terminal GST fusions in *E.coli* BL21(DE3) and purified using Glutathione Sepharose 4B (Cytiva). Briefly, recombinant protein expression was induced with 0.5 mM IPTG (isopropyl-thio-β-D-galactopyranoside) at 18°C overnight. Cell pellets were resuspended in PBS containing 0.5 mM TCEP (tris(2-carboxyethyl) phosphine) and lysed by sonication. GST fusion proteins were affinity-purified from soluble bacterial lysates using Glutathione Sepharose 4B (Cytiva) according to standard manufacturer’s protocols. After elution in 25 mM reduced glutathione, 50 mM Tris pH7.4, 350 mM NaCl, 0.5 mM TCEP, recombinant proteins were rebuffered into PBS, 0.5 mM TCEP using PD-10 desalting columns (GE Healthcare) followed by concentration with Amicon Ultra centrifugation filter devices (Millipore) and stored at −80°C until required. E2 enzymes were purified analogously. UBE1 was purified as previously described^11^. Cy5-labeled ubiquitin was prepared as previously described^36^.

Full-length ZNFX1 was recombinantly expressed in insect or human cells. Briefly, for expression in insect cells, pOP806_pACEBac1 2×Strep-ZNFX1 (wild-type) plasmid was transformed into DH10EmBacY cells. Blue–white screening was used to isolate colonies containing recombinant baculoviral shuttle vectors (bacmids) and bacmid DNA was extracted combining cell lysis and neutralization using buffer P1, P2 and N3 (Qiagen), followed by isopropanol precipitation. A 6-well plate of *Spodoptera frugiperda* (Sf9) cells (Oxford Expression Technologies) grown at 27°C in Insect-Xpress media (Lonza) without shaking was transfected with bacmid using PEI transfection reagent. After 7 days, virus P1 was collected and used 1:20 to transduce 100 mL (2.0 × 10^6^ cells/ml) of Sf9 cells. After 7 days of incubation at 27°C with 140 rpm shaking, virus P2 was collected. To express protein, 4 L (2.0 × 10^6^ cells/mL) of Sf9 cells were transduced with 1:40 dilution of P2 virus and incubated at 27°C with 140 rpm shaking for 72 hours. Cells were pelleted by centrifugation, snap-frozen in liquid nitrogen, and stored at −80°C. To lyse cells, the pellets were thawed and resuspended to 200 mL total volume in lysis buffer (30 mM HEPES, pH 7.6, 100 mM NaCl, 10 mM MgCl_2_, 0.5 mM TCEP), containing 20 μL universal nuclease (Pierce), 20 μL benzonase nuclease (Sigma Aldrich), and 4 EDTA-free protease inhibitor tablets (Roche). The cell suspension was stirred for 1 hour and then sonicated for 50 seconds in 5-second pulses with 25-second waiting time at 70% amplitude using a 130 W microtip sonicator (Vibra Cell). The lysate was centrifuged at 20,000×g for 20 min at 4°C. The clarified lysate was filtered through a 5.0 μm filter (Millipore), topped up to increase the salt concentration to 300 mM NaCl, and applied to 2 × 5-mL StrepTrap HP columns (GE Healthcare), connected in series. After washing the columns with 100 mL lysis buffer, supplemented with NaCl in a linear gradient (from 300 mM to 500 mM and back down to 100 mM NaCl), ZNFX1 was eluted with lysis buffer, supplemented with 2.5 to 10 mM desthiobiotin, pH 8. The eluted protein was kept at 4°C and used immediately for cryo-EM specimen preparation.

ZNFX1 (wild type and mutants) was expressed in adherent HEK293T *ZNFX1* KO cells for biochemical assays. A 10 cm plate of cells at around 80–90% confluency was transfected with 12 µg of plasmid (Peak8-Strep-HA-GFP-ZNFX1) using PEI. After 24 hours cells were harvested in PBS and lysed in buffer A (50 mM Tris-HCl, pH8, 150 mM NaCl, 2 mM MgCl2), supplemented with 1% NP40, 1:1000 benzonase nuclease, 1:500 BioLock, 1×protease inhibitor cocktail) for 10 minutes on ice. The lysates were clarified by centrifugation at 20,000×g for 5 minutes. StrepTactin 4Flow beads were added to the clarified lysates and incubated for 1 hour at 4°C. The beads were washed with 3 × 1 mL high-salt wash buffer (buffer A + 1% NP40 + 350 mM NaCl), then 3 × 1 mL wash buffer (buffer A + 1% NP40), then 2 × 1 mL buffer A. Elution buffer (buffer A + 10 mM desthiobiotin) was added to the resin, which was then transferred to a spin column, and the eluate was collected and used immediately for biochemical assays.

### In vitro ubiquitylation assays

In vitro ubiquitylation experiments were set up in reaction buffer (30 mM HEPES pH7.4, 100 mM NaCl, 10 mM MgCl2, 5 mM ATP) containing 100 nM E1 (UBE1), 200 nM E2, 10 μM Cy5-labeled ubiquitin and 300-500 nM GST-ZNFX1 fragments or full-length ZNFX1 protein. Reactions were incubated at 37°C for 60 min, stopped by adding 2x sample buffer, separated on SDS-PAGE, and then analyzed by in-gel fluorescence and Coomassie staining.

### Cryo-EM specimen preparation

Recombinant ZNFX1 was purified as described above. For the first dataset (ZNFX1 + ATP), ZNFX1 was used at a concentration of 3.0 mg/mL and ATP was added to a final concentration of 0.5 mM. For the second dataset (ZNFX1 + RNA), a single-stranded 200-nucleotide RNA molecule was added at 50 ng/uL to ZNFX1 at 1.0 mg/mL concentration. The mixed samples were incubated at room temperature for 10 minutes and kept on ice afterwards. All-gold grids^37^ (UltrAuFoil R 1.2/1.3, 300-mesh, QuantiFoil Micro Tools GmbH) were used for all specimens. The grids were plasma cleaned in an atmosphere of 9:1 Ar:O_2_ for 2 minutes at 70% power, 30 sccm gas flow in a Fischione 1070 plasma chamber. The samples (3 µL per grid) were vitrified using a Vitrobot Mark IV (ThermoFisher Scientific), operated at 4°C and 100% relative humidity with 5-second blotting time and blot force 10. The grid was then immediately plunged into liquid ethane, held at 93 Kelvin in a temperature-controlled cryostat^38^. All grids were stored in liquid nitrogen until use.

### Cryo-EM data collection

Cryo-EM data were collected on a Titan Krios (ThermoFisher Scientific) electron cryomicroscope, operated at 300 keV and equipped with a Selectris X (ThermoFisher Scientific) imaging filter with a post-filter Falcon 4i (ThermoFisher Scientific) direct electron detector. Both datasets were acquired using EPU (ThermoFisher Scientific) with aberration-free image shift within a 6 μm range and correspond to 24 hours of data collection each. Data collection parameters are summarized in **Supplementary Table S1**.

### Cryo-EM data processing

Both datasets were processed using a combination of RELION-5.0^39^ and cryoSPARC-4.5^40^. Briefly, patch-based motion correction, CTF estimation, and initial particle blob-picking were performed in cryoSPARC Live for both datasets.

For the first dataset, (ZNFX1 + ATP), the picked particles were extracted into 360-pixel boxes and binned by a factor of 3.6. The extracted particles were subjected to multiple rounds of 2D classification. Selected 2D classes were used for template-based re-picking and the newly picked particles were extracted into 500-pixel boxes and binned by a factor of 5. The particles were subjected to multiple rounds of 2D classification, ab-initio reconstruction with multiple classes, and heterogeneous refinement, where false picks and damaged particles were removed. The selected particles yielded an initial reconstruction of the ZNFX1 filament. A mask was created around the repeating unit in the filament (a dimer of two ZNFX1 molecules with C2 symmetry), and further local refinement was performed with this mask and with C2 symmetry applied. Finally, the selected particles were re-extracted into 500-pixel boxes and re-binned into 450-pixel boxes. The optical aberrations were estimated in cryoSPARC. Finally, local 3D refinement with regularization by the Blush algorithm, was performed in RELION-5.0^41^, followed by C2-symmetry expansion and a final round of local refinement with Blush regularization using a mask around a single ZNFX1 monomer.

For the second dataset (ZNFX1 + RNA), the picked particles were extracted into 400-pixel boxes and binned by a factor of 2. The particles were subjected to multiple rounds of 2D classification, ab-initio reconstruction with multiple classes, and heterogeneous refinement, where false picks and damaged particles were removed. Finally, the selected particles were re-extracted into 400-pixel boxes without binning, and exported into RELION, where the optical aberrations were estimated. After performing local 3D refinement with regularization by the Blush algorithm, 3D classification without alignment was performed using a small mask around the RNA-binding groove in the helicase domain to select for the RNA-bound molecules. The final particle set was once again subjected to local 3D refinement with Blush regularization.

The efficiency of the orientation distributions for both datasets was calculated using cryoEF^42^. Graphics were generated in ChimeraX^43^.

### Atomic model building and refinement

The initial atomic model was generated by piece-wise docking of fragments of the structure of human ZNFX1, as predicted using AlphaFold 3.0^44^ into the cryo-EM map. The model was manually refined in ISOLDE^45^ and in Coot 0.9.4 with Geman-McClure restraints (α=0.003)^46^, followed by real-space refinement in PHENIX^47^ with secondary structure restraints and Ramachandran restraints.

### Multiple sequence alignment

Pair-wise sequence alignments were performed using Clustal Omega^48^; all other multiple sequence alignments were performed using MUSCLE^49^. ZNFX1 homologs from multiple species were retrieved from the Ensembl database^50^ using the REST API^51^, and only sequences longer than 1,500 amino acids (an arbitrarily chosen cut-off intended to remove poor quality, truncated sequences) were retained for alignment. Conservation scores were calculated using the ConSurf Server^52^.

## Acknowledgements

This work was supported by the Medical Research Council as part of United Kingdom Research and Innovation (U105170648, to FR) and the Wellcome Trust (222503/Z/21/Z, to FR), (217191/Z/19/Z, to YM). We acknowledge Diamond for access and support of the cryo-EM facilities at eBIC, proposal BI31336, funded by the Wellcome Trust, MRC and BBSRC. MY was supported by an EMBO Postdoctoral Fellowship. We thank Lori Passmore for helpful discussions. We also thank the MRC LMB cryo-EM, scientific computing, light microscopy, insect cell culture, mass spectrometry, and flow cytometry facilities for their support.

## Data availability

The cryo-EM maps and the refined atomic models have been deposited in Electron Microscopy Data Bank and the Protein Data Bank. Other data is available within the article and its associated supplementary information files.

## Competing interests

The authors declare no competing financial interests.

## Author contributions

KB, JH, CH, AH, KL, TM, KN, EGO and MY performed experiments and analyzed the results. VL, YM and FR provided funding for this study. TM, KN, MY and FR wrote the manuscript with input from all authors.

## Supplementary Figures

**Figure S1.**
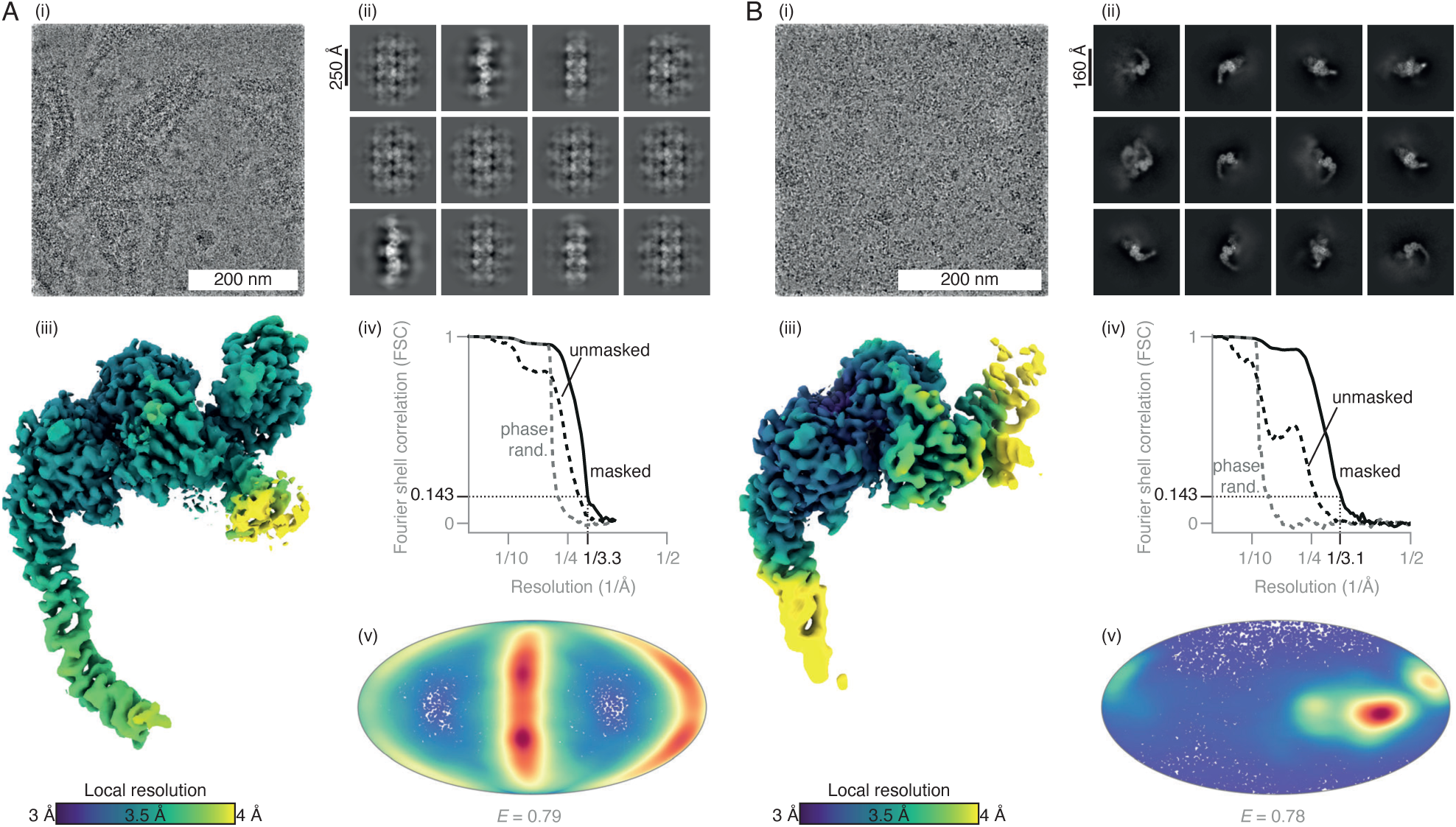
Cryo-EM structure determination of human ZNFX1 in complex with ATP and in complex with RNA. (A) Cryo-EM data processing for the ZNFX1 + ATP dataset. (i) Representative micrograph at 2.4 µm defocus, low-pass filtered to 1 nm resolution for display. (ii) Selected 2D class averages, showing filamentous ZNFX1, and bundles of these filaments. (iii) Reconstruction of ZNFX1 bound to ATP, colored by local resolution. (iv) Fourier shell correlation as function of resolution for the final masked map (black), the unmasked map (dashed black), and the phase-randomized map (dashed grey). (v) Orientation distribution, with efficiency *E*, of the particles used in the final reconstruction. (B) Cryo-EM data processing for the ZNFX1 + RNA dataset. (i) Representative micrograph at 2.9 µm defocus, low-pass filtered to 1 nm resolution for display. (ii) Selected 2D class averages, showing monomeric ZNFX1. (iii) Reconstruction of ZNFX1 bound to RNA, colored by local resolution. (iv) Fourier shell correlation as function of resolution for the final masked map (black), the unmasked map (dashed black), and the phase-randomized map (dashed grey). (v) Orientation distribution, with efficiency *E*, of the particles used in the final reconstruction.

**Figure S2.**
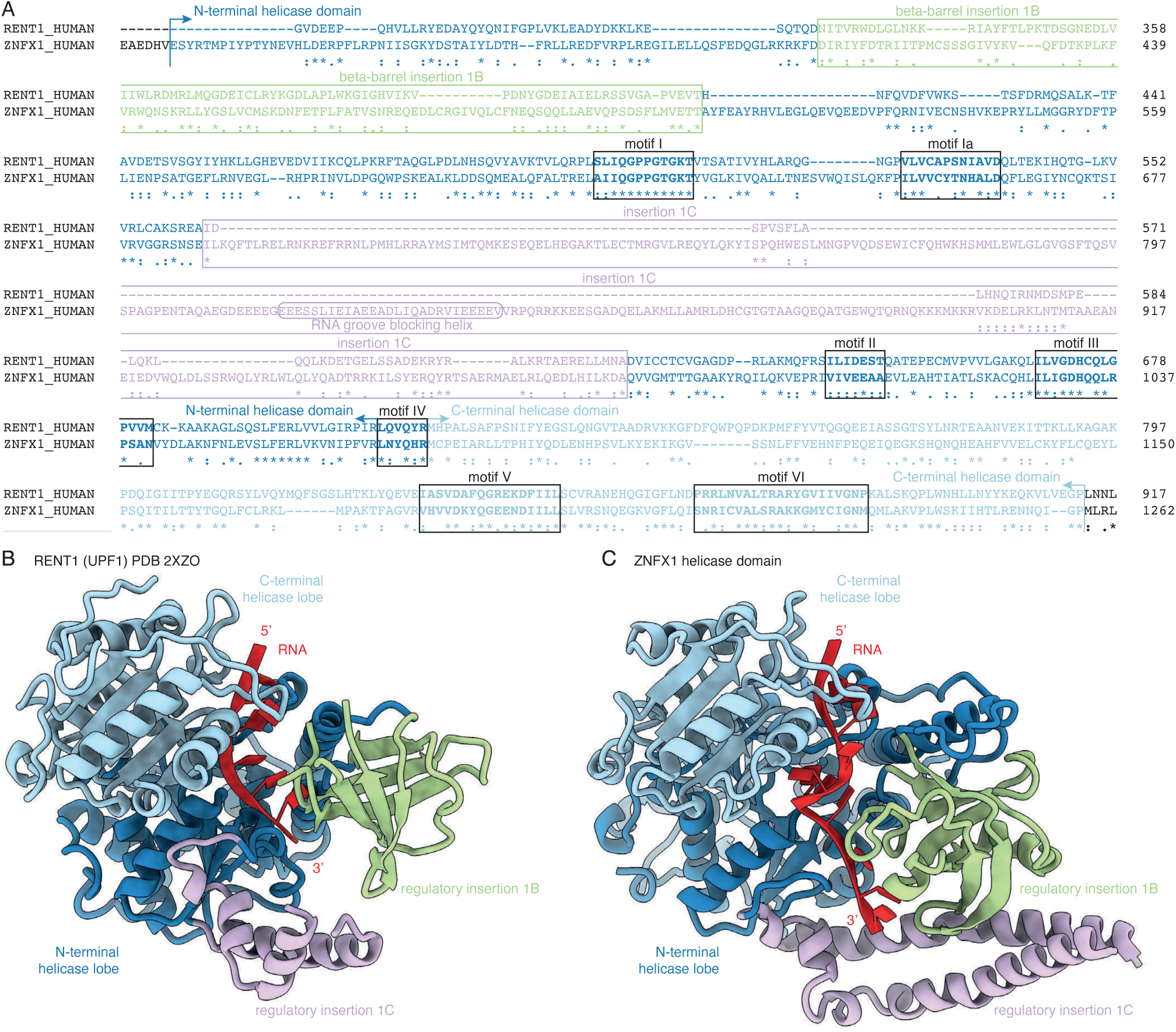
Comparison between the RNA helicase domains of human ZNFX1 and human Upf1 (RENT1). (A) Protein sequence alignment between human Upf1 and human ZNFX1. The seven motifs, typical of SF1-family helicases, are boxed. The sub-domains are colored as in Figure 1. (B) Structure comparison between the Upf1 (PDB 2XZO)^16^ and ZNFX1 helicase domains, in the RNA-bound state.

**Figure S3.**
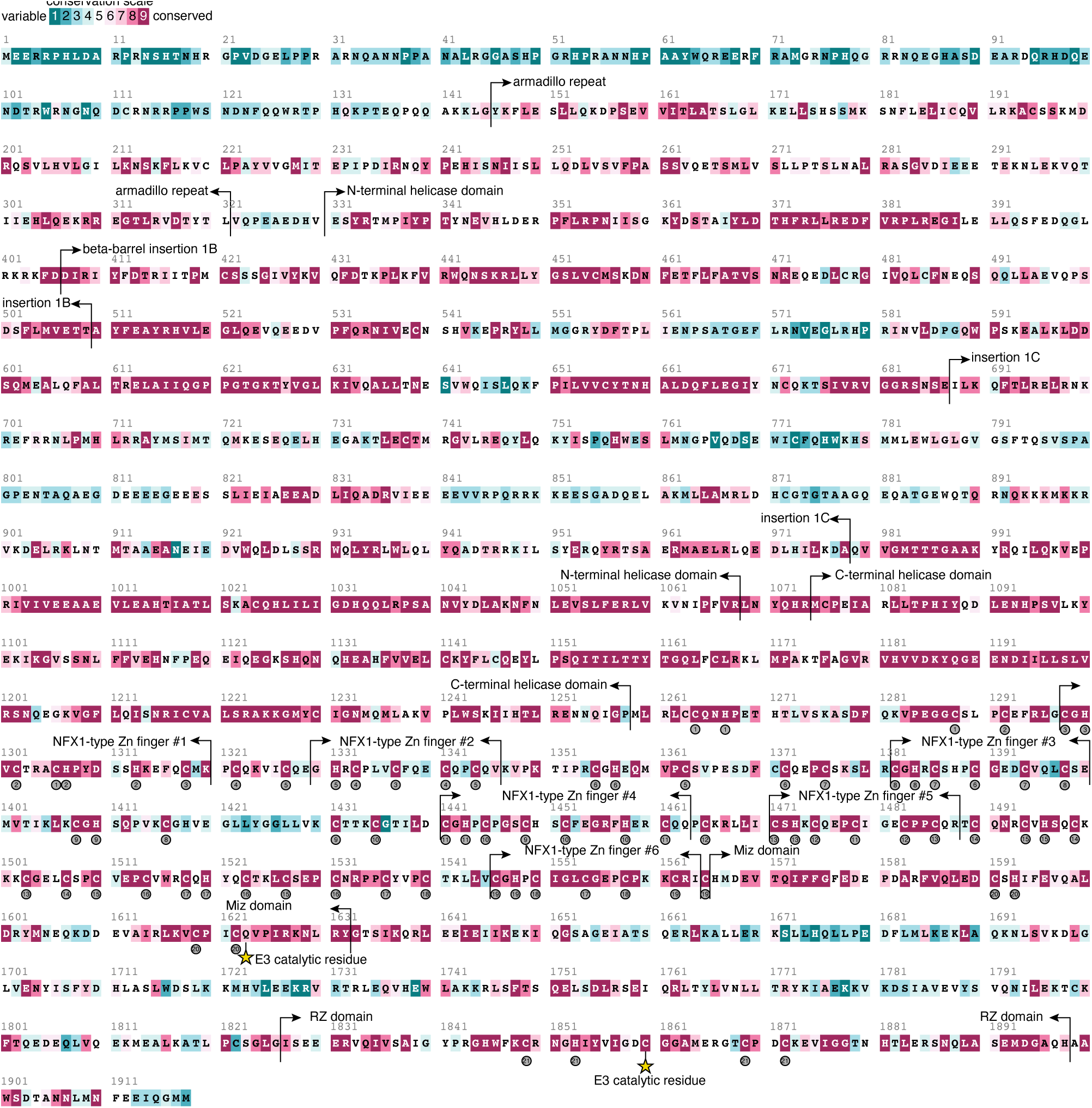
ZNFX1 conservation across species. The human ZNFX1 protein sequence is colored by conservation, based on a sequence alignment of ZNFX1 orthologs from a total of 185 species, and is annotated by domains. The numbers in grey circles refer to the predicted Zn coordination sites.

**Figure S4.**
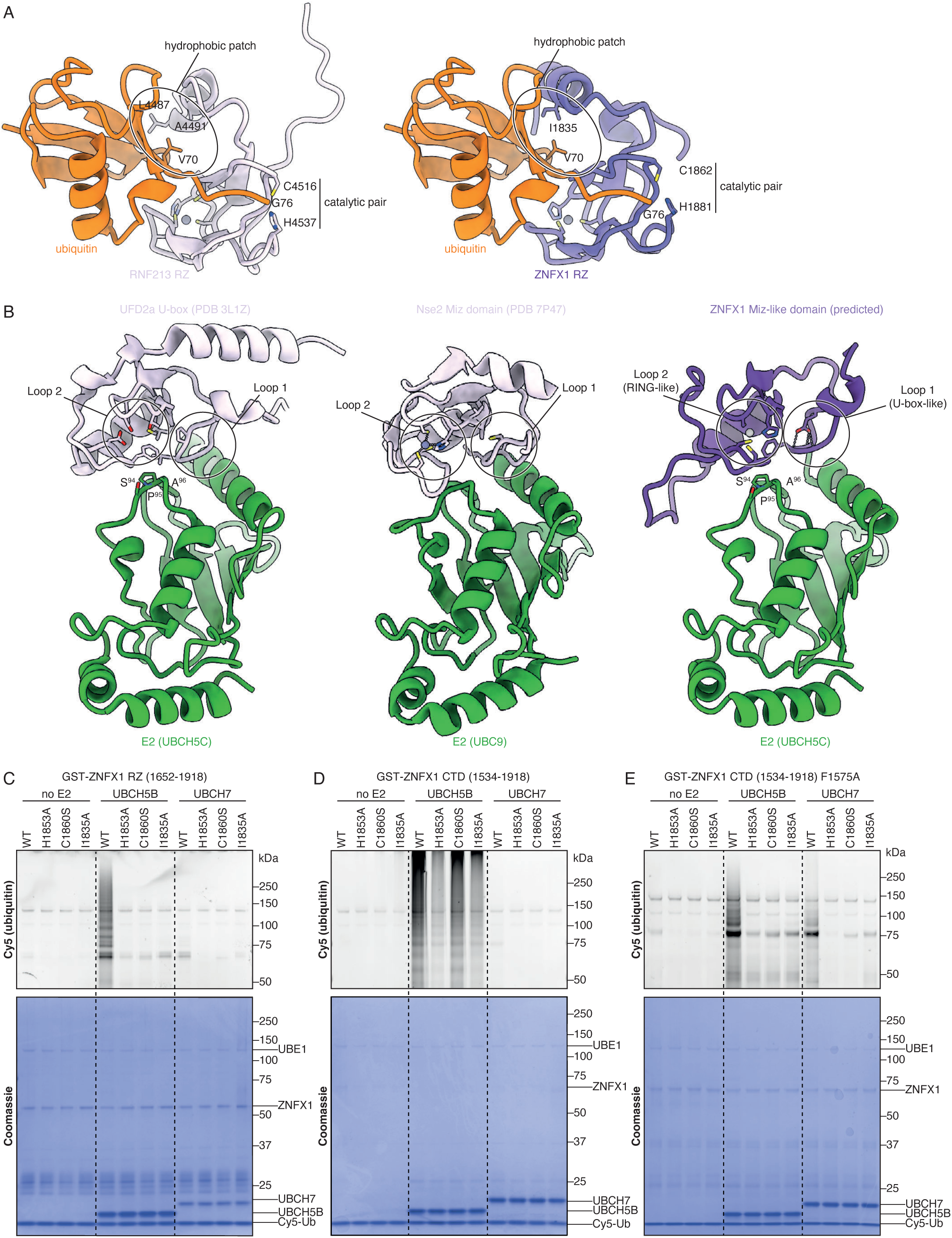
ZNFX1 E3 ubiquitin ligase domains: structural analogues and *in* vitro activity. (A) Comparison between the predicted structures of RNF213 RZ and ZNFX1 RZ domains bound to ubiquitin. (B) Comparison between the previously determined structures of the UFD2 U-box domain, bound to E2 enzyme from the UBCH5C family, PDB 3L1Z^53^, the Nse2 Miz domain, bound to the SUMO-specific E2 enzyme UBC9, PDB 7P47^54^, and the predicted structure of the ZNFX1 Miz-like domain in complex with UBCH5C. (C) E3 ubiquitin ligase catalytic activity of the ZNFX1 RZ domain (wild type and mutants) with the UBCH5B, UBCH7, or no E2 enzyme control, monitored *in vitro* by Cy5-ubiquitin fluorescence and Coomassie staining of SDS-PAGE (full figure of Figure 4B). (D) Same as in (C), but for the ZNFX1 C-terminal domain (CTD), containing both the Miz domain and the RZ domain. (E) Same as in (D), but for the ZNFX1 CTD containing a mutation (F1575A) that disrupts E2 binding to the Miz domain. All ZNFX1 fragments for (C– E) were expressed in bacteria.

**Figure S5.**
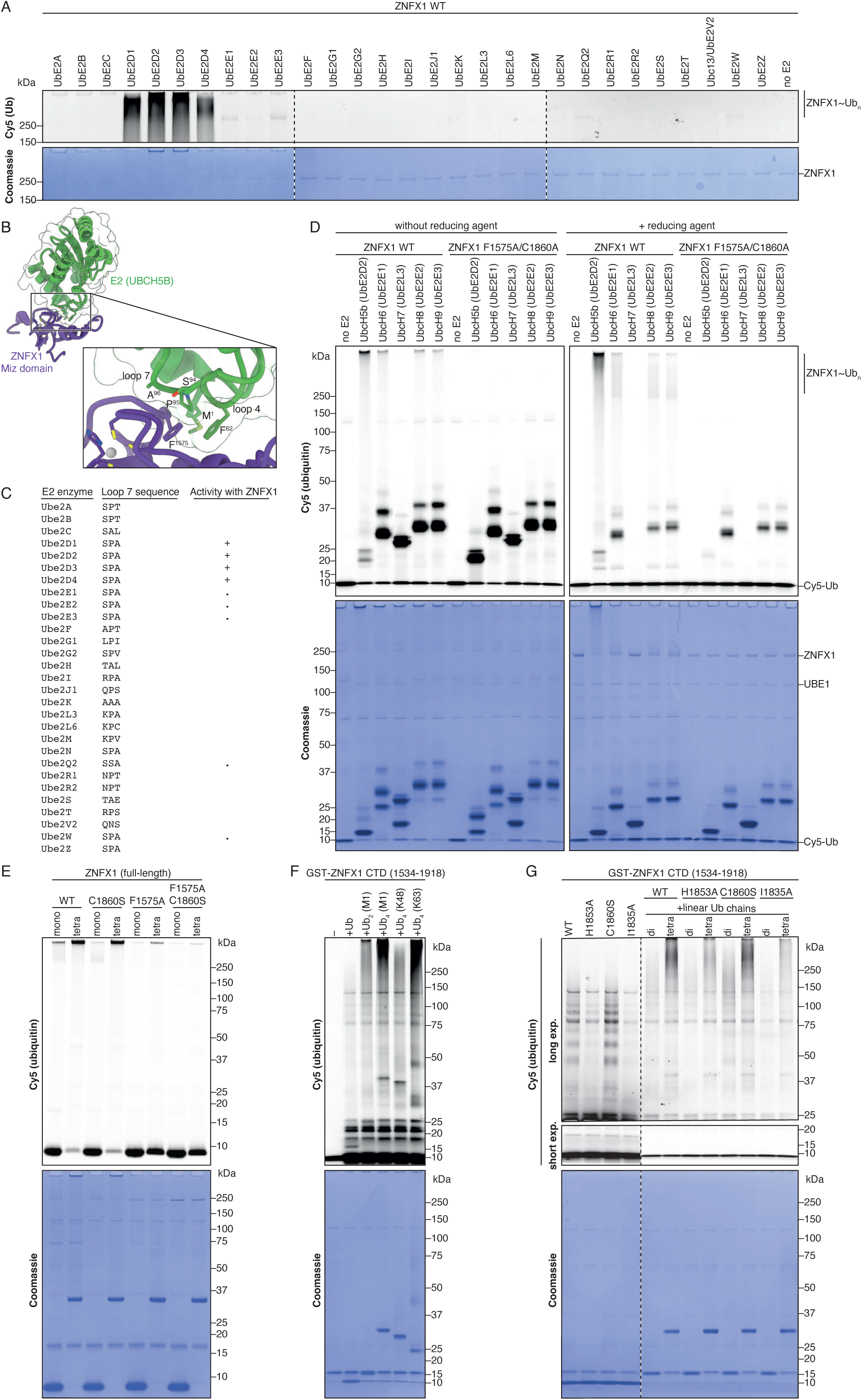
E2 enzyme selectivity and allosteric activation of ZNFX1. (A) *In vitro* ubiquitylation assays with full-length ZNFX1, purified from human cells, and a panel of different human E2 enzymes (UBPBio), where activity is monitored by Cy5-ubiquitin fluorescence and Coomassie staining of SDS-PAGE. (B) Predicted structure of the Miz-type domain of ZNFX1 in complex with UBCH5B, highlighting residues forming the interaction interface in the inset. (C) Sequence alignment of the typical E3-binding region (loop 7) of all human E2 enzymes, and their activity with ZNFX1 (“+” indicates good activity, “.” indicates low activity). (D) Same as in (A) but comparing wild-type and E2 binding/RZ inactive double mutant (F1575A/C1860A) ZNFX1 in the presence of the indicated E2 enzymes. The assays were monitored *in vitro* by Cy5-ubiquitin fluorescence and Coomassie staining of SDS-PAGE. (E) *In vitro* ubiquitylation assay with the indicated variants of full-length ZNFX1, in the presence of additional unlabeled mono-ubiquitin or linear (M1-linked) tetra-ubiquitin chains. (F) *In vitro* ubiquitylation assay with a wild-type ZNFX1 C-terminal domain (CTD) fragment, purified from bacteria, in the presence of additional unlabeled mono-ubiquitin, linear (M1-linked) di-or tetra-ubiquitin, and K48-or K63-linked tetra-ubiquitin. (G) *In vitro* ubiquitylation assay with the indicated variants of the ZNFX1 CTD fragments, purified from bacteria, in the absence (left) or presence (right) presence of additional unlabeled linear (M1-linked) di-or tetra-ubiquitin. In all assays (E–G) the amount of Cy5-labeled ubiquitin is the same across all conditions and corresponds to approximately 1 part in 20 of all ubiquitin monomers in the reaction.

## Supplementary Tables

**Supplementary Table S1.**
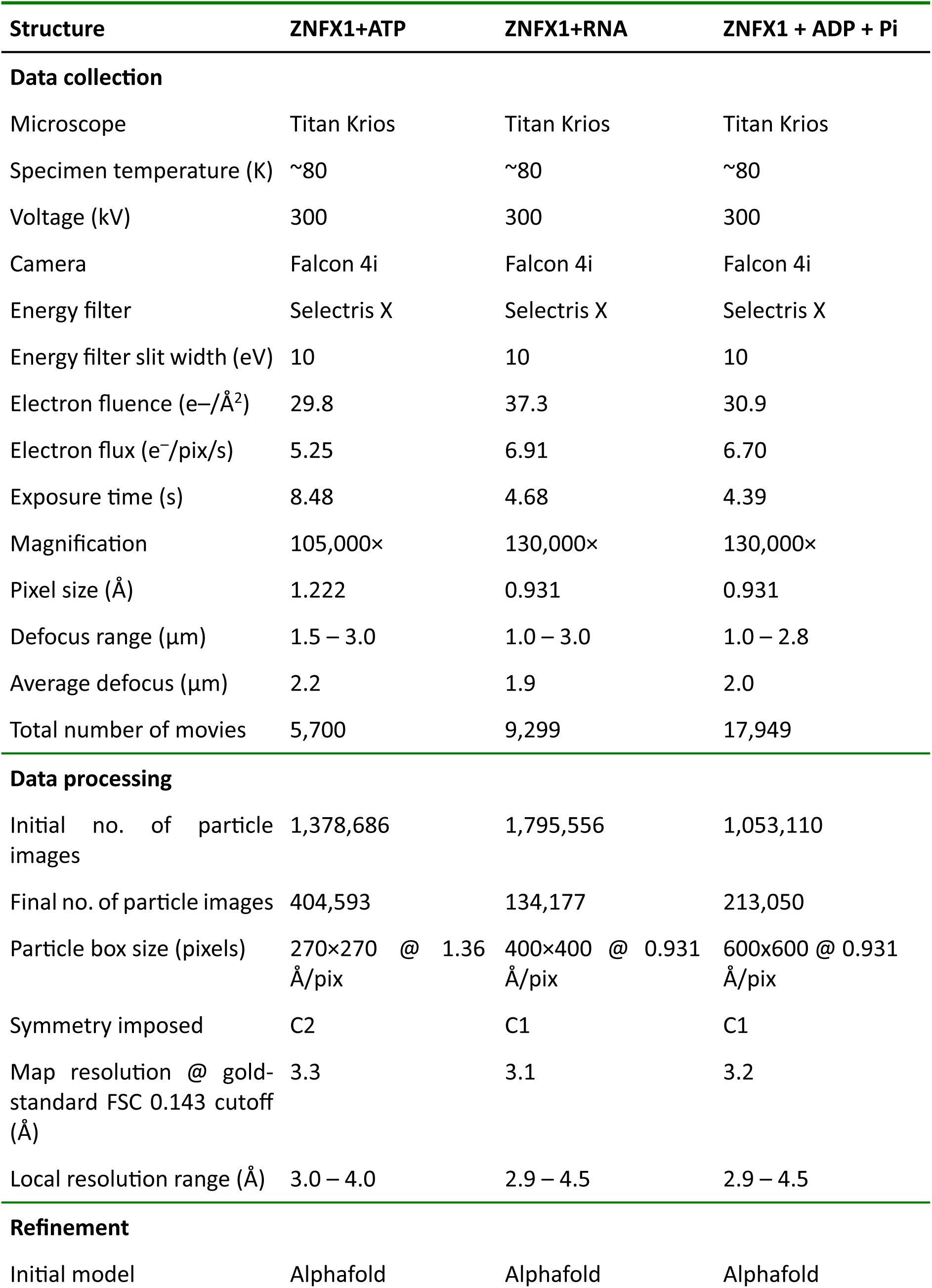
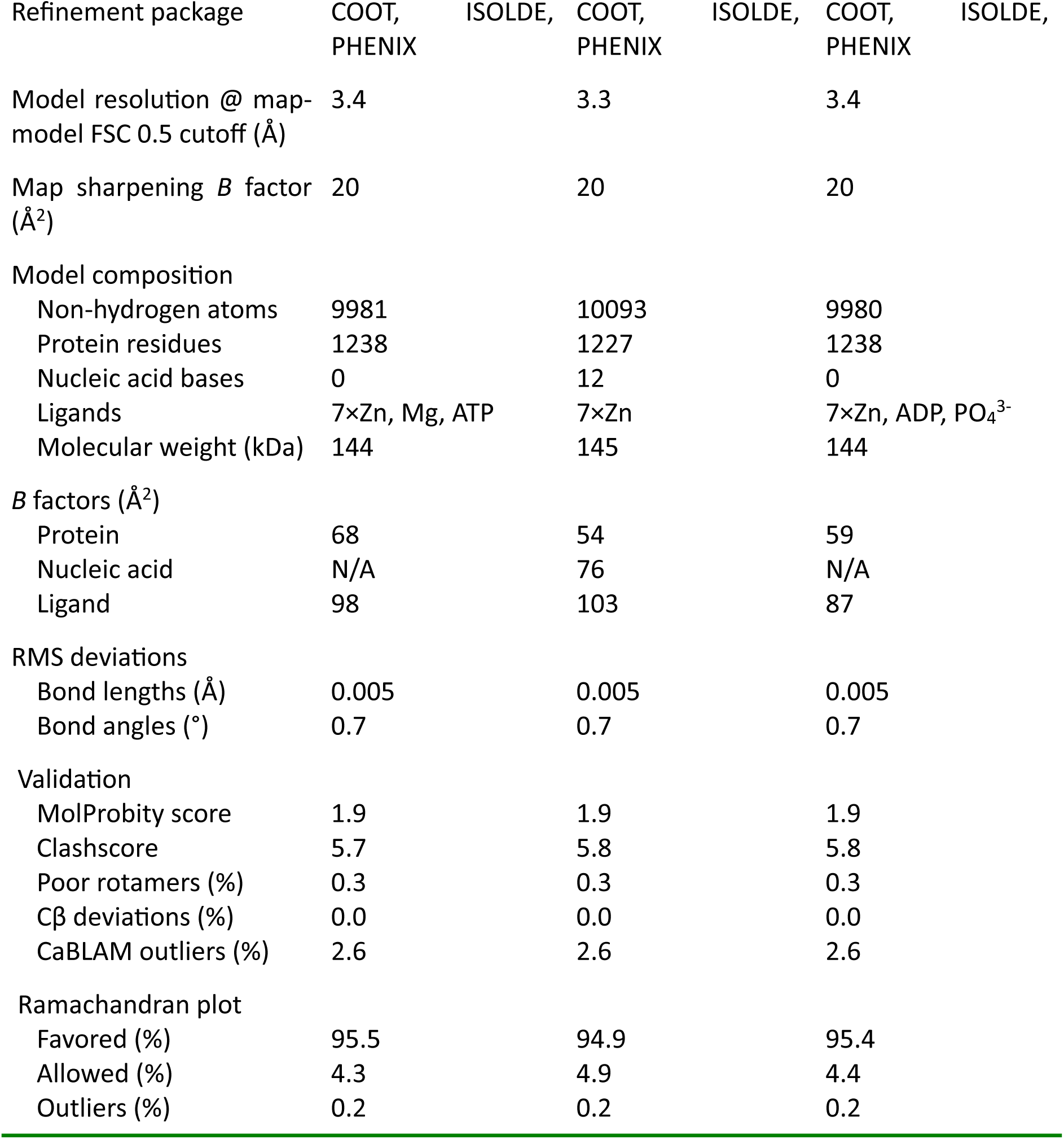
Cryo-EM data collection and processing, model refinement and validation statistics.

**Supplementary Table S2.** Previously reported mutations in *ZNFX1* of clinical significance. Any mutations that result in truncation of the predicted protein product before the end of the helicase domain have been omitted from this list, as they most likely either result in nonsense-mediated decay of the corresponding short transcripts or in a truncated ZNFX1 protein fragment completely devoid of any of its catalytic activities.

| Mutation | Domain | Reference |
| --- | --- | --- |
| V469L | Beta-barrel insertion 1B in N-terminal helicase lobe (conserved residue) | <sup>5</sup> |
| L1051P | N-terminal helicase lobe | <sup>4</sup> |
| R1223Q | C-terminal helicase lobe (conserved residue) | <sup>5</sup> |
| C1264S; E1727Kfs*11 | Zn fingers (coordinating residue for Zn ion #1); terminates within C-terminal helical bundle with complete loss of RZ domain | <sup>4</sup> |
| I1154T; C1292S | C-terminal helicase lobe; Zn fingers (coordinating residue for Zn ion #2) | <sup>4</sup> |
| E1606Rfs*10 | Terminates within U-box with loss of Zn finger | <sup>3</sup> |
| V1788Cfs*6 | C-terminal helix bundle, terminates with loss of Zn finger | <sup>5</sup> |

## Notes

### Competing Interest Statement

The authors have declared no competing interest.

